# Receptor binding domain of SARS-CoV-2 is a functional αv-integrin agonist

**DOI:** 10.1101/2022.04.11.487882

**Authors:** Emma G. Norris, Xuan Sabrina Pan, Denise C. Hocking

**Author notes:** **To whom correspondence should be addressed:** Denise C. Hocking, Ph.D., Box 711, 601 Elmwood Ave., University of Rochester, Rochester, NY 14642, USA.

## Abstract

Among the novel mutations distinguishing SARS-CoV-2 from similar respiratory coronaviruses is a K403R substitution in the receptor-binding domain (RBD) of the viral spike (S) protein within its S1 region. This amino acid substitution occurs near the angiotensin-converting enzyme 2 (ACE2)-binding interface and gives rise to a canonical RGD adhesion motif that is often found in native extracellular matrix proteins, including fibronectin. In the present study, the ability of recombinant S1-RBD to bind to cell surface integrins and trigger downstream signaling pathways was assessed and compared to RGD-containing, integrin-binding fragments of fibronectin. S1-RBD supported adhesion of both fibronectin-null mouse embryonic fibroblasts as well as primary human small airway epithelial cells. Cell adhesion to S1-RBD was cation- and RGD-dependent, and was inhibited by blocking antibodies against α_v_ and β_3_, but not α_5_ or β_1_, integrins. Similarly, direct binding of S1-RBD to recombinant human α_v_β_3_ and α_v_β_6_ integrins, but not α_5_β_1_ integrins, was observed by surface plasmon resonance. Adhesion to S1-RBD initiated cell spreading, focal adhesion formation, and actin stress fiber organization to a similar extent as fibronectin. Moreover, S1-RBD stimulated tyrosine phosphorylation of the adhesion mediators FAK, Src, and paxillin, Akt activation, and supported cell proliferation. Together, these data demonstrate that the RGD sequence within S1-RBD can function as an α_v_-selective integrin agonist. This study provides evidence that cell surface α_v_-containing integrins can respond functionally to spike protein and raise the possibility that S1-mediated dysregulation of ECM dynamics may contribute to the pathogenesis and/or post-acute sequelae of SARS-CoV-2 infection.

## Introduction

Coronaviruses (CoVs) are a diverse group of positive-stranded RNA viruses named for the distinctive crown-like protrusions on their surfaces. CoVs can infect a wide range of mammalian and avian species, causing mild to severe respiratory infections, and were first identified as human pathogens in the 1960s (1). At present, 7 different CoVs are known to infect humans, 4 of which cause only mild disease (2). Within the past 20 years, 3 CoVs have emerged that are capable of causing more severe disease in humans: SARS-CoV, the cause of Severe Acute Respiratory Syndrome (SARS), MERS-CoV, the cause of Middle East Respiratory Syndrome (MERS), and SARS-CoV-2, the cause of COVID-19 (1). Since its emergence in Wuhan, China in late 2019, there have been 446 million reported cases, leading to over 6 million deaths (3). The most common symptoms of infection are fever, cough, shortness of breath, and fatigue (4), but disease progression varies widely with approximately 20% of non-vaccinated patients experiencing severe acute disease (5). In patients presenting with severe disease, common risk factors include advanced age, diabetes, hypertension and other cardiovascular diseases (4). Acute respiratory distress syndrome (6), as well as myocardial (7), renal (8), hepatic (9),and digestive (10) complications have all been reported. Additionally, over half of COVID-19 patients, including those with mild, acute symptoms, exhibit a range of short and long-term post-acute sequelae that include pulmonary abnormalities, functional mobility impairment, fatigue, and joint pain (11,12). The complex clinical manifestations of acute and post-acute COVID-19 suggest a dysregulated host response to infection that triggers immuno-inflammatory, thrombotic and parenchymal disorders (13), Yet, the pathophysiological mechanisms responsible for the diverse disease phenotypes remain largely unknown.

The extracellular matrix (ECM) glycoprotein, fibronectin is an essential regulator of connective tissue homeostasis (14), epithelial morphogenesis (15), endothelial barrier maintenance (16,17), local arteriolar tone (18,19), and tissue repair (20). Fibronectin also serves a significant role in host-pathogen interactions, as fibronectin-binding and fibronectin-mimicking proteins have been identified across a broad spectrum of microbial pathogens (21). Compared to SARS-CoV, the spike (S)1 subunit of SARS-CoV-2 contains a novel mutation that mimics a bioactive sequence in fibronectin: a Lys (K) to Arg (R) mutation in the receptor binding domain (RBD), resulting in the adhesive Arg-Gly-Asp (RGD) motif of fibronectin’s integrin-binding domain (22). In fibronectin, the RGD sequence is located in a short loop that extends from the tenth type III repeat (FNIII10) where it mediates adhesion for a variety of cell types, including epithelial cells, endothelial cells and fibroblasts, via β_1_ and β_3_ integrins (23,24). Ligation of cell surface integrins with the RGD sequence of fibronectin triggers a cascade of cell signaling events, including protein kinase C activation and Rho-mediated actomyosin contractility, that lead to changes in cell shape (25), focal adhesion composition (26,27), and extracellular matrix assembly (28). Critically, activation of components of these adhesion-based signaling cascades has been associated with reduced endothelial and epithelial barrier function and increased prevalence of inflammatory diseases (29).

SARS-CoV-2 infects epithelial cells of both the respiratory (30) and gastrointestinal (GI) tracts (31) via S1-mediated recognition of angiotensin-converting enzyme 2 (ACE2) on host cell surfaces (32). Initial evaluation of inter-residue distances within the crystal structure of SARS-CoV-2 spike in complex with ACE2 suggested that the RGD motif of S1 is located adjacent to, but not included within, the ACE2-binding surface (33). More recent analysis indicates that Arg403 of S1 is highly conserved across SARS-CoV-2 lineages, and may facilitate viral engagement of human cells via an ionic interaction with residue Glu37 of ACE2 (34). Positive detection of S1-integrin binding via solid-phase ELISA assays has been reported by several independent groups for both α_5_β_1_ (35,36) and α_v_β_3_ (37) integrins. Viral infection studies further showed that cell-surface binding and viral uptake of SARS-CoV-2 can be inhibited by integrin antagonists, including the peptide inhibitors Cilengitide (37) and ATN-161 (35,38), as well as by cell-permeable inhibitors of inside-out integrin signaling (39). Thus, converging evidence suggests that S1-integrin interactions occur during SARS-CoV-2 infection, though the specificity and selectivity for specific integrins, as well as the implications for SARS-CoV-2 infection and disease remain to be elucidated. In the present study, we investigated S1-integrin interactions using both primary human small airway epithelial cells, as well as fibronectin-null mouse embryonic fibroblasts (FN-null MEFs). FN-null MEFs do not produce fibronectin, laminin, or vitronectin (40,41) and are cultured in the absence of serum, allowing for the characterization of S1 binding to cell surface receptors and the identification of intracellular signals triggered by S1-integrin engagement without interference from other adhesive ligands (41–43). Results of this study indicate that the RGD motif contained within S1 is a cryptic, low-affinity, α_v_ integrin ligand that can mediate cell adhesion, spreading and proliferation to a similar extend as native fibronectin. RBD-integrin engagement triggers canonical integrin-mediated signaling cascades, focal adhesion formation, and actin cytoskeletal organization, thus functioning as a classical α_v_ integrin agonist.

## Results

### S1-RBD of SARS-CoV2 supports cell adhesion and proliferation via α_v_β_3_ integrins

The integrin-binding RGD motif is contained within a variety of endogenous ECM glycoproteins (44) and is frequently expressed by microbial pathogens as a mechanism for attachment to host tissue (45). To begin to determine whether S1-RBD functionally interacts with cells, FN-null MEFs were seeded into wells coated with either S1-RBD or the RGD-containing module of fibronectin, FNIII10. At 4 h after seeding, cells adherent to S1-RBD exhibited robust adhesion and classical fibroblast morphology, characterized by extended membrane protrusions (Fig. 1A). Cell adhesion was dose-dependent with respect to substrate coating concentration, and comparable to adhesion on FNIII10 (Fig. 1B). Interestingly, full-length S1 supported only minimal cell adhesion compared to the similarly-sized fibronectin fragment, FNIII8-13 (Fig. 1C), suggesting that the integrin-binding epitope may not be accessible in the larger S1 fragment.

**Figure 1.**
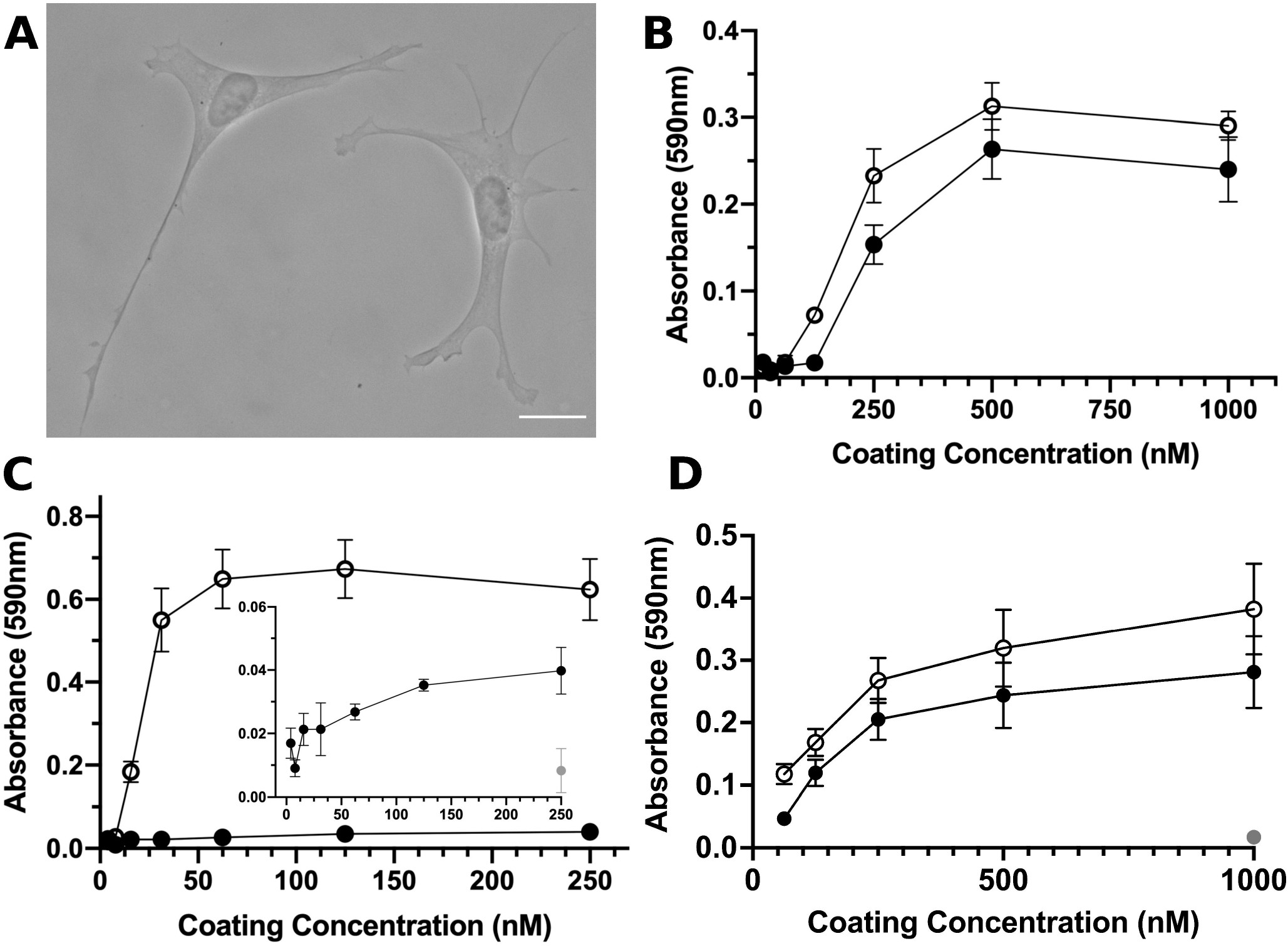
S1-RBD supports cell adhesion and proliferation. **A**, FN-null MEFs (2.5×10 cells/cm) were seeded onto coverslips pre-coated with S1-RBD (500 nM) and cultured for 4 h prior to fixation and phase-contrast imaging. Scale bar, 20 μm. **B**, FN-null MEFs (1.9×10^5^ cells/cm^2^) were seeded onto tissue culture plates precoated with the indicated concentration of S1-RBD (filled circles) or FNIII10 (open circles). Cells were cultured for 90 min and relative cell number was determined by crystal violet staining. **C**, FN-null MEFs (1.9×10^5^ cells/cm^2^) were seeded onto plates pre-coated with HN-tagged S1 (filled circles) or FNIII8-13 (open circles) for 90 min. Inset shows cell adhesion to S1 (filled circles) compared to BSA-coated wells (gray circle). **D**, FN-null MEFs (2.3×10^3^ cells/cm^2^) were seeded onto tissue culture plates pre-coated with the indicated concentration of S1-RBD (filled circles), FNIII10 (open circles), or GST (gray circle) and cultured for 4 d. Relative cell number was determined by crystal violet staining. Data are mean ± SEM; n ≥ 3 experiments performed in triplicate.

Ligation of integrins by RGD-containing agonists initiates cell signaling cascades that support cell proliferation (46). Thus, we next tested the ability of S1-RBD to support cell proliferation. To do so, FN-null MEFs were seeded at low density in defined, serum-free media onto tissue culture plates coated with S1-RBD, FNIII10 or the nonadhesive, protein purification tag, glutathione S-transferase (GST). After a 4-day incubation, relative cell number was quantified as a function of coating concentration. As shown in Fig. 1D, cell number increased similarly with increasing coating density on S1-RBD- and FNIII10-coated wells. In contrast, cells seeded into GST-coated wells did not survive (Fig. 1D; 1 μM), indicating that cell proliferation in response to S1-RBD was specific and not due to the presence of endogenous or exogenously-supplied adhesive proteins.

Integrins are heterodimeric receptors, whose ligand specificity is determined by the combination of alpha and beta subunits (44). FN-null MEFs express α_1_, α_5_, α_v_, β_1_, and β_3_ integrin subunits (41), of which both α_5_β_1_ and α_v_β_3_ are RGD-binding integrins (44). To identify the integrin receptors mediating cell adhesion to S1-RBD, FN-null MEFs were pre-incubated with blocking antibodies directed against β_1_, β_3_, α_5_, or α_v_ integrin subunits. Cell adhesion to S1-RBD was inhibited partially by antibodies against β_3_-integrin subunits, and inhibited completely by either a combination of α_v_- and β_3_-blocking antibodies, or EDTA (Fig. 2A). Similar results were obtained using the α_v_β_3_-specific ligand, FNIII10 (47) (Fig. 2B). In contrast, cell adhesion to S1-RBD was not inhibited by either α_5_- or β_1_-blocking antibodies (Fig. 2A) under conditions that specifically inhibited adhesion to the β_1_-integrin ligand, collagen I (Fig. 2C). Rather, treatment of cells with anti-β_1_ antibodies significantly increased cell adhesion to S1-RBD (Fig. 2A). Finally, competitive inhibition assays were performed using short RGD-, RAD-, or KGD-containing peptides. Addition of soluble RGD peptides blocked cell adhesion to S1-RBD (Fig. 2D). In contrast, addition of control, RAD peptides had no effect on cell adhesion to S1-RBD (Fig. 2D). Furthermore, peptides derived from the RGD-containing region of SARS-CoV-2 partially inhibited cell adhesion to S1-RBD, whereas a peptide derived from the corresponding sequence of SARS-CoV-1, which contains a KGD rather than RGD motif, did not reduce adhesion to S1-RBD (Fig. 2D). Notably, SARS-CoV-2-derived peptides also partially inhibited cell adhesion to FNIII10 (Fig. 2E). Together, these data indicate that the RGD motif of S1-RBD ligates α_v_β_3_ integrins in a cationdependent manner.

**Figure 2.**
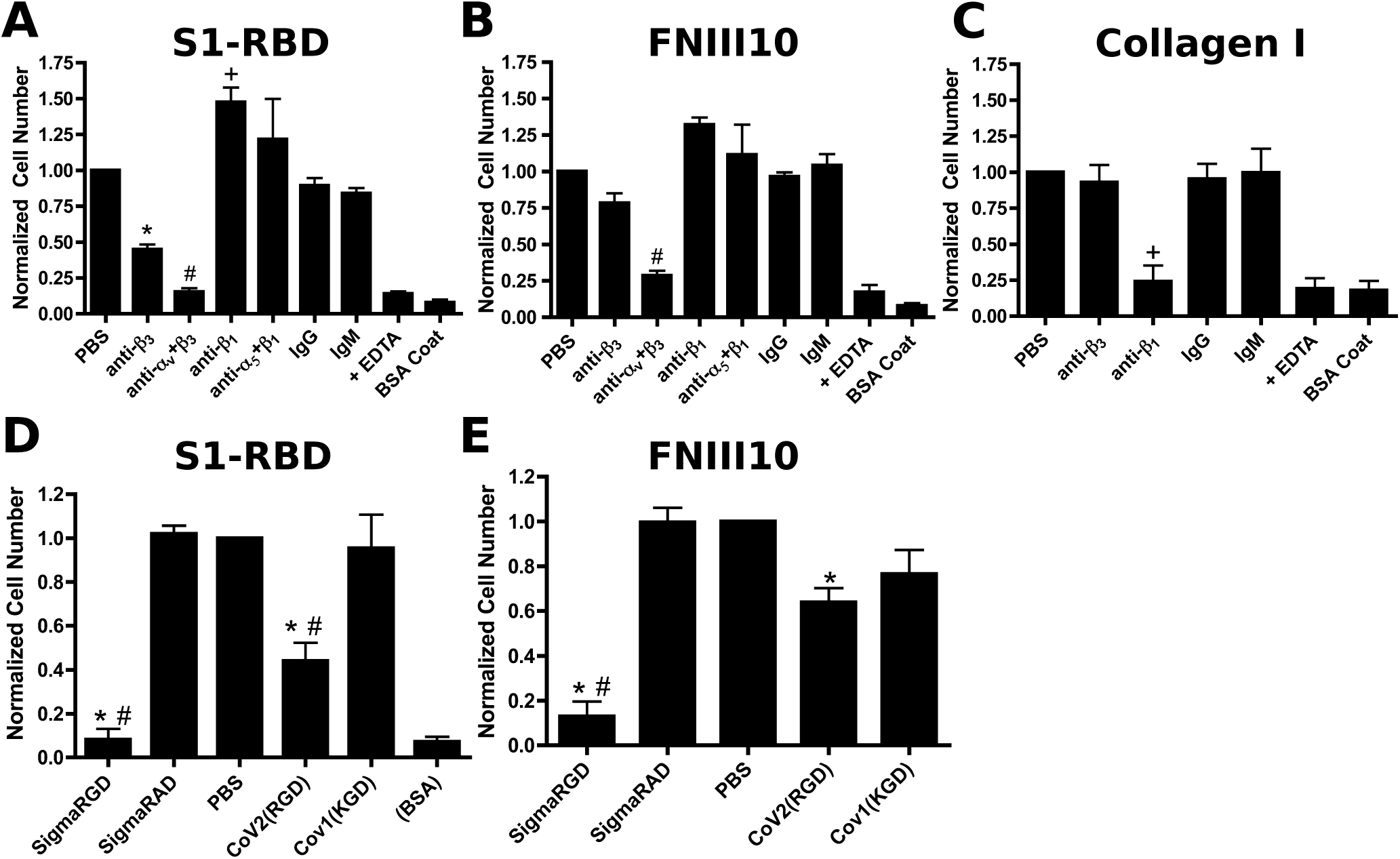
FN-null MEF adhesion to S1-RBD is mediated by α_v_β_3_ integrins and RGD. FN-null MEFs (5 x 10^5^ cells/mL) were pre-incubated for 30 min with 50 μg/mL integrin-blocking antibodies (**A-C**) or 25 μM peptide (**D-E**) before seeding (9.4 x 10^4^ cells/cm^2^) onto plates pre-coated with 250 nM S1-RBD (**A,D**), FNIII10 (**B,E**), or type I collagen (**C**). Relative cell number was determined by crystal violet staining. Data are mean ± SEM, normalized to corresponding vehicle (PBS) controls; n=3 independent experiments performed in triplicate. One-way ANOVA, Bonferroni’s post-hoc test: **A-C** *p<0.05 vs PBS, IgG; +p<0.05 vs PBS, IgM; #p<0.05 vs PBS, anti-α_5_+β_1_; **D-E** *p<0.05 vs PBS; #p<0.05 vs corresponding negative control, RAD or KGD.

Surface plasmon resonance (SPR) was used next to study the kinetic parameters governing the binding of recombinant human integrins with immobilized S1-RBD. Representative response curves obtained from α_v_β_3_, α_v_β_6_ or α_5_β_1_ integrin binding to S1-RBD, FNIII10, or FNIII8-10 are shown in Fig. 3. Experimental data were collected and globally fit using a 1:1 binding model. The fitted kinetic parameters, k_a_, k_d_, K_D_, and Rmax are shown in Table 1. Quality of each fit was evaluated by comparison to the experimentally measured Rmax and to Chi^2^. Measurable binding of α_v_β_3_ integrins to S1-RBD was observed (Fig. 3A). However, curve-fitting parameters indicated that the goodness of fit was not sufficient to perform kinetic analysis, suggesting that the K_D_ of α_v_β_3_ integrins binding to S1-RBD is greater than 500 nM (Table 1). Kinetic modeling of data obtained for α_v_β_6_ integrin binding to S1-RBD provided a K_D_ value of 230 nM (Fig. 3B, Table 1). In contrast, α_5_β_1_ integrins did not bind to S1-RBD (Fig. 3C), in agreement with results of cell adhesion assays (Fig. 2). Measured affinities of FNIII10 binding to α_v_β_3_ (Fig. 3D) and α_v_β_6_ (Fig. 3E) integrins were 21.8 nM and 6.6 nM, respectively (Table 1), which are similar to published values (48). Association rates for the interaction of α_v_β_6_ integrins with S1-RBD and FNIII10 were similar (S1-RBD k_a_ = 8.4 x 10^4^ M^-1^s^-1^; FNIII10 k_a_ = 7.1 x 10^4^ M^-1^s^-1^). In contrast, the dissociation rate of α_v_β_6_ integrins with S1-RBD was much larger than that observed with FNIII10 (S1-RBD k_d_ = 236 x 10^-4^ s^-1^; FNIII10 k_d_ = 6.0 x10^-4^ s^-1^). Kinetic fits were not performed for the reaction of S1-RBD and α_5_β_1_ integrins, as binding was not observed (Fig. 3C) even at analyte concentrations substantially exceeding the K_D_ of the interaction of FNIII8-10 with α_5_β_1_ integrins (48) and under reaction conditions in which α_5_β_1_ integrin binding to FNIII8-10 was observed (Fig. 3F). Together, these data indicate that S1-RBD is capable of binding directly to α_v_ integrins through low affinity interactions.

**Figure 3.**
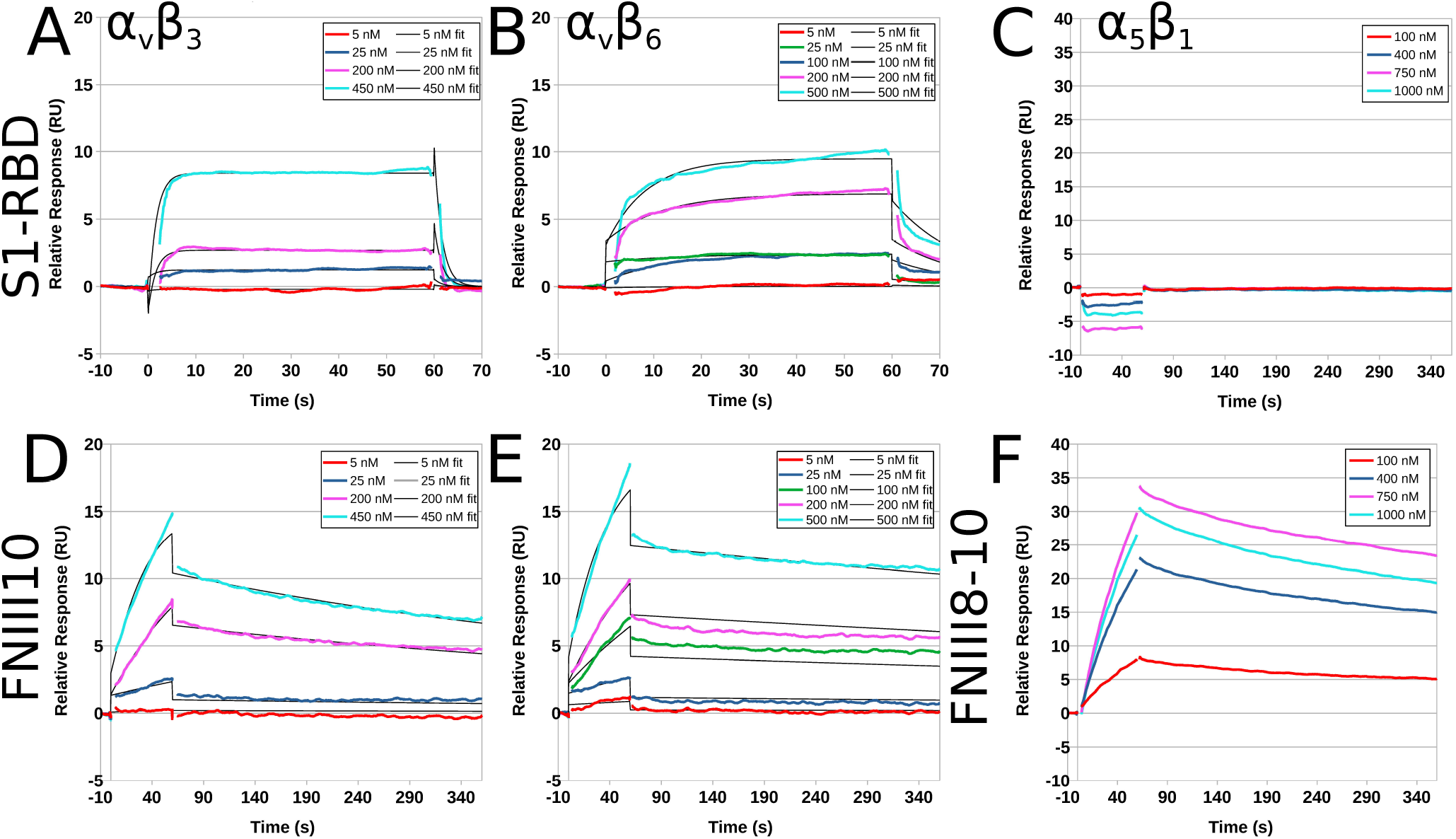
Recombinant human integrins α_v_β_3_ and α_v_β_6_, but not α_5_β_1_, bind to immobilized S1-RBD. Representative kinetic data for α_v_β_3_ (**A, D**), α_v_β_6_ (**B, E**) or α_5_β_1_ (**C, F**) binding to immobilized S1-RBD (**A-C**), FNIII10 (**D-E**), or FNIII8-10 (**F**). Data are presented as representative traces (colored lines) collected from 1 of 2 (α_5_β_1_) or 3 (α_v_β_3_ and α_v_β_6_) experiments and corresponding 1:1 binding fits (black lines).

**Table 1.** Summary of kinetic and quality control parameters determined for the interaction of α_v_ integrins with immobilized S1-RBD and FNIII10. Data are presented as mean ± SEM for at least 3 independent experiments per integrin. Double-referenced experiments were performed simultaneously for S1-RBD and FNIII10 ligands in parallel flow cells.

| Ligand | Analyte | $k_a \times 10^4$ | $k_d \times 10^{-4}$ | $K_D$ (nM) | $R_{max}$ fit (RU) | $R_{max}$ measured (RU) | $\chi^2$ |
| --- | --- | --- | --- | --- | --- | --- | --- |
| S1-RBD | $\alpha_v\beta_3$ | - | - | > 500 | - | - | - |
| S1-RBD | $\alpha_v\beta_6$ | $8.4 \pm 1.8$ | $236 \pm 206$ | $230 \pm 180$ | $6.9 \pm 3.5$ | $11.7 \pm 0.7$ | $0.1 \pm 0.3$ |
| FNIII10 | $\alpha_v\beta_3$ | $7.1 \pm 1.4$ | $15.0 \pm 1.8$ | $21.8 \pm 1.6$ | $12.8 \pm 0.5$ | $12.6 \pm 0.4$ | $0.06 \pm 0.01$ |
| FNIII10 | $\alpha_v\beta_6$ | $25.0 \pm 18.8$ | $6.0 \pm 0.7$ | $6.6 \pm 3.1$ | $16.6 \pm 2.5$ | $19.0 \pm 2.3$ | $0.16 \pm 0.05$ |

### S1-RBD initiates focal adhesion formation and actin organization

Integrin ligation by endogenous ECM ligands triggers adhesion signaling cascades in which intracellular mediators are recruited to sites of integrin activation (46,49). These protein complexes, known as focal adhesions, serve as central signaling hubs and functionally couple ECM-engaged integrins to the actin cytoskeleton (50). Notably, manipulation of focal adhesion signaling has been identified across a diverse spectrum of microbial pathogens, with the potential to influence multiple stages of cellular pathophysiology, including cell-surface attachment, invasion, and cell death (45). To determine whether engagement of α_v_β_3_ integrins by S1-RBD supports focal adhesion formation and downstream signaling, FN-null MEFs adherent to S1-RBD were stained with the actin-binding protein phalloidin, together with antibodies against the focal adhesion adaptor vinculin, and a pan-specific phosphotyrosine antibody (51). Cells seeded on S1-RBD- or FNIII10-coated substrates exhibited classical features of focal adhesions, including co-localized vinculin and phosphotyrosine staining, as well as actin stress fiber formation (Fig. 4).

**Figure 4.**
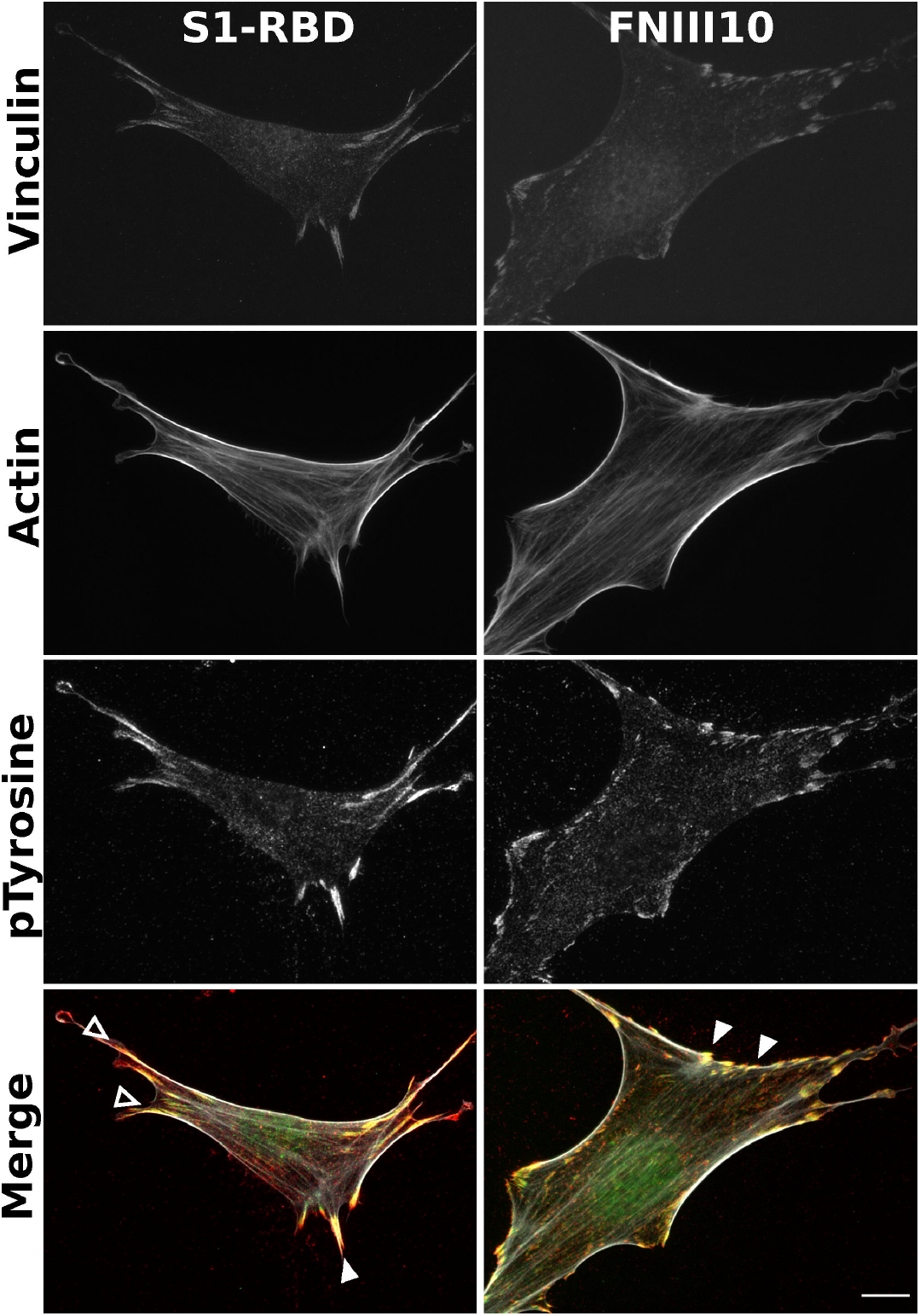
S1-RBD engagement initiates focal adhesion formation and actin organization. FN-/- MEFs (2.5×10 cells/cm^2^) were seeded on coverslips coated with 500 nM S1-RBD (left) or FNIII10 (right). Cells were incubated for 4 h prior to fixation and immunofluorescent staining for vinculin (green), actin (TRITC-phalloidin, white), or phospho-tyrosine (4G10, red). Arrowheads represent co-localization of vinculin and phosphotyrosine within focal adhesions (closed) and engagement with the actin cytoskeletion (open). Representative images shown from 1 of 4 independent experiments. Scale bar, 10 μm.

To identify proteins specifically phosphorylated by S1-RBD ligation, immunoblot analysis of whole-cell lysates were performed. Similar patterns of protein tyrosine phosphorylation were observed when lysates from attached S1-RBD- or FNIII10-adherent cells were probed with a pan-specific phosphotyrosine antibody (not shown). As such, immunoblots were next probed with phosphospecific antibodies against key components of adhesion signaling pathways (Fig. 5). These components included the early, adhesion-dependent autophosphorylation of focal adhesion kinase (FAK) at Y^397^(52), which in turn enables recruitment and phosphorylation of Src at Y^418^ (53). Both FAK-Y^397^ and Src-Y^418^ were phosphorylated in response to S1-RBD ligation (Fig. 5). Moreover, the extent of FAK and Src phosphorylation was similar to that observed in either FNIII-10- or fibronectin-adherent cells (Fig. 5). FAK may be phosphorylated at additional tyrosine residues including Y^407^(54), which was phosphorylated to a similar extent in both suspended and adherent cells (Fig. 5). S1-RBD triggered tyrosine phosphorylation of paxillin (Fig. 5), a central adaptor protein whose SH2 domains require phosphorylation at residues Y^118^ and Y^31^ for activation and cytoskeletal remodeling (55,56). Additionally, S1-RBD induced Akt phosphorylation at residue S^473^ to a similar extent as FNIII10- and fibronectin-adherent cells, implicating engagement of the pro-survival PI3K/Akt signaling axis (57,58) and consistent with results demonstrating that S1-RBD ligation supports cell proliferation (Fig. 1D). Together, these data indicate that S1-RBD can trigger multiple aspects of adhesion-based signaling, including localization of vinculin to focal adhesions, phosphorylation of early adhesion signals FAK, and Src, as well as activation of downstream adhesive effectors, including paxillin and Akt.

**Figure 5.**
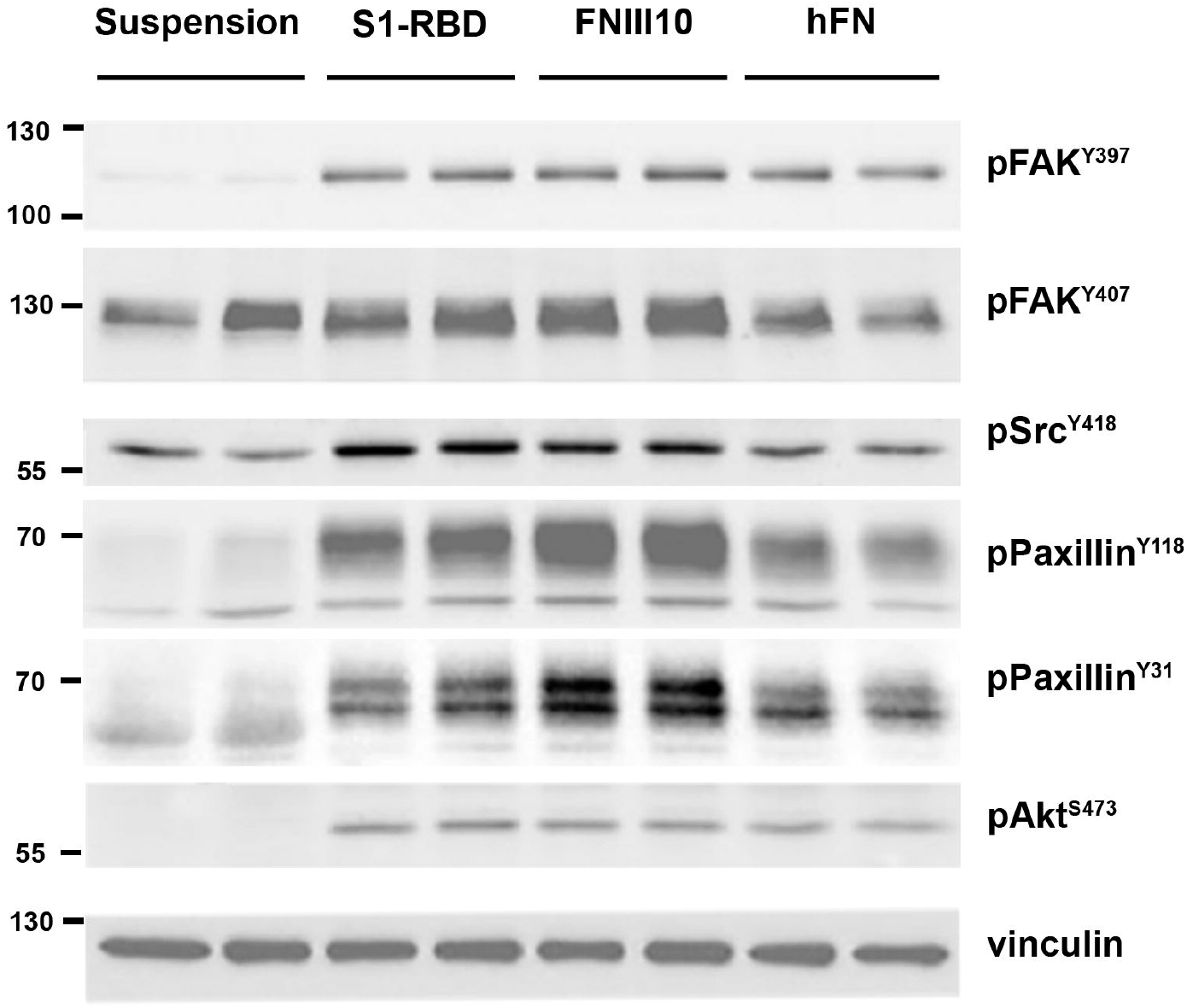
Cell engagement with S1-RBD stimulates intracellular signaling. FN-null MEFs were either suspended in media or seeded at 3.5×10^5^ cells/cm^2^ on wells pre-coated with 500 nM of S1-RBD or FNIII10, or 10 μg/ml of human plasma fibronectin (hFN) for 1 h. Cell lysates were analyzed by immunoblotting with the indicated phosphospecific antibodies. Molecular mass markers are shown at left

Similar adhesion assays and immunoblot analyses were performed using human small airway epithelial cells (hSAECs). These primary cells are derived from the distal lung, and are susceptible to SARS-CoV-2 infection (59,60). S1-RBD supported hSAEC adhesion to a similar extent as FNIII10 (Fig. 6A). Furthermore, hSAECs adhesion to S1-RBD stimulated tyrosine phosphorylation of FAK, paxillin, and Src to a similar extent as FNIII10 and fibronectin (Fig. 6B). Thus, these data indicate that S1-RBD stimulates adhesion-mediated intracellular signaling pathways in human lung epithelial cells.

**Figure 6.**
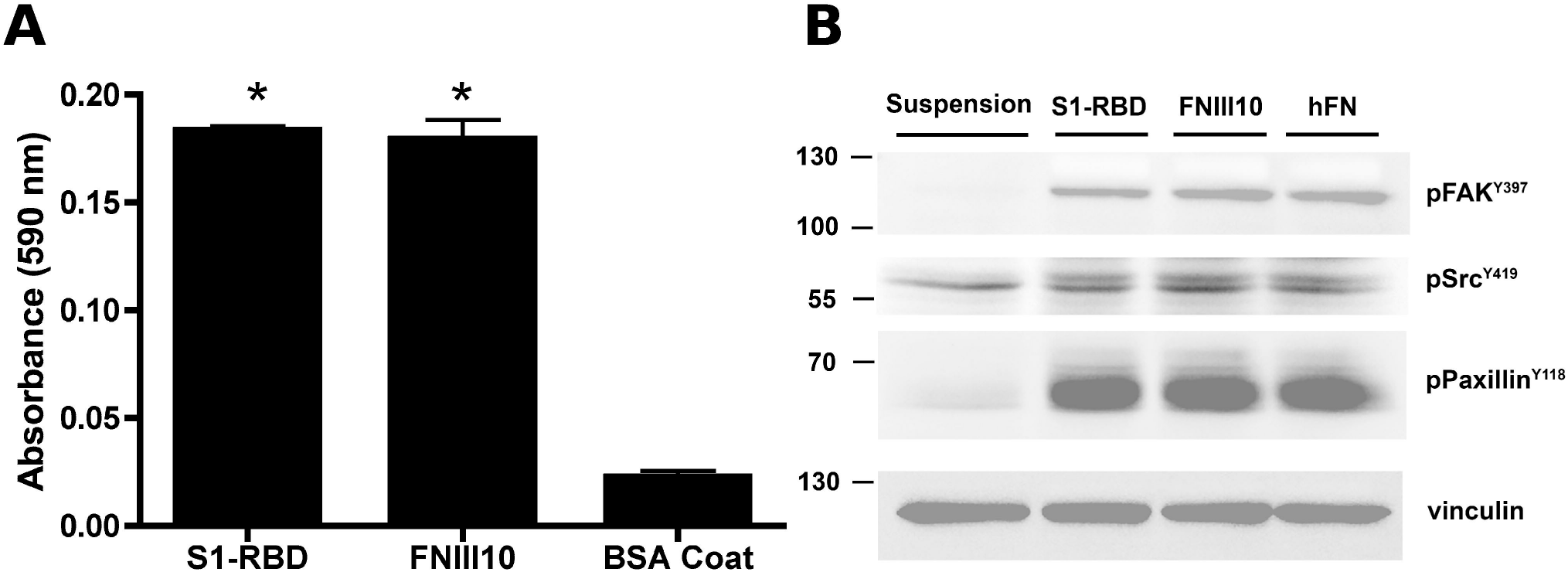
S1-RBD supports cell adhesion and phosphotyrosine signaling in human small airway epithelial cells. hSAECs (6.7 x 10^4^ cells/cm^2^) were seeded in the presence of 1 mM DTT on wells pre-coated with S1-RBD, FNIII10, or 1% BSA. **A**, Cell adhesion was determined by crystal violet staining after 2 h. Data are presented as mean absorbance +/- SEM for n=3 independent experiments. *P<0.05 vs BSA by one-way ANOVA with Bonferroni’s post-test. **B**, Cell lysates were obtained after 4 h of adhesion and analyzed by immunoblotting with the indicated phosphospecific antibodies. Molecular mass markers are shown at left. Control cells were maintained suspended in media before lysing.

### Adhesive epitope contained within S1 spike protein is conformationally sensitive

The data presented thus far indicate that the RGD sequence within S1-RBD is a functional, integrin-binding ligand that can mimic classical features and functions of native ECM ligands, including cation-sensitive adhesion, cell spreading, focal adhesion formation, phosphotyrosine signaling, and cell proliferation. In contrast, cells attached poorly to a larger fragment of S1 (Fig. 1C), which contains both the N-terminal domain and furin cleavage site in addition to RBD. Cryptic adhesive epitopes are a feature common to many native ECM proteins, including fibronectin (61) and thrombospondin (62). To determine whether the adhesive activity of S1 is cryptic, we chemically reduced recombinant S1 protein by treatment with dithiothreitol (DTT). Reduced S1, but not control (non-reduced) S1, supported a significant increase in cell adhesion compared to non-adhesive BSA (Fig. 7A). Notably, cell adhesion to reduced S1 protein also required the presence of MnCl_2_, which potently activates both α_v_β_3_ and α_5_β_1_ integrins via binding to their metal ion-dependent adhesion site domain (63,64). The Mn^+2^-dependence of cell adhesion to S1 is in contrast to cell adhesion to the S1-RBD fragment, which supported cell adhesion in both the presence and absence of MnCl_2_ (Fig. 7A). A small, but not statistically significant increase was observed in the number of cells attached to non-reduced S1 in the presence of MnCl_2_ (Fig. 7A). Cells adherent to non-reduced S1 exhibited a rounded morphology with minimal cell spreading (Fig. 7B). In contrast, cell spreading was observed on reduced S1 (Fig. 7C). Cells did not attach to non-adhesive BSA, even in the presence of MnCl_2_ (Fig. 7D). Saturating cell adhesion, with robust cell spreading was observed for both the S1-RBD (Fig. 7E) and FNIII10 (not shown) cell fragments, with no differences observed in the presence or absence of MnCl_2_. Together, these data indicate that the RGD motif of RBD is cryptic within the S1 fragment, and that this site can be exposed by disulfide reduction and integrin activation.

**Figure 7.**
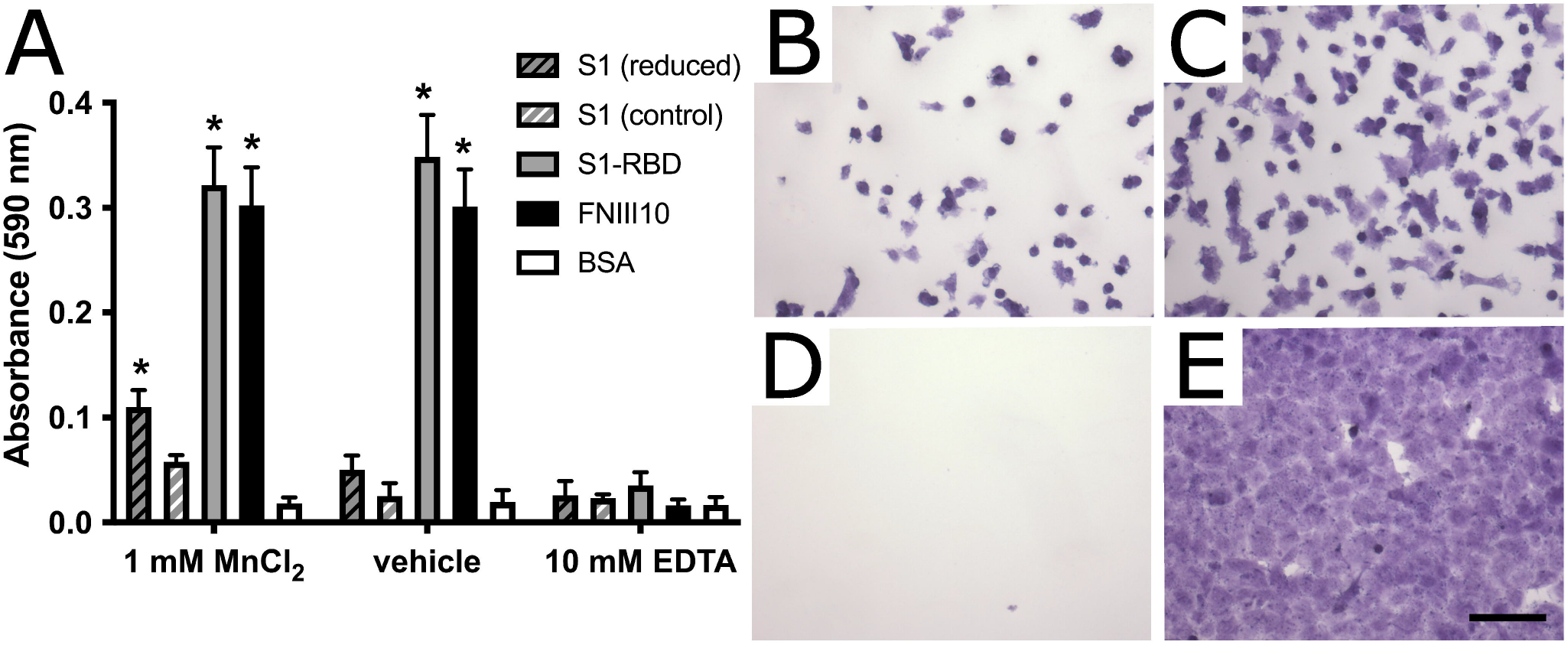
S1 contains a cryptic, DTT- and Mn^2+^-sensitive adhesive epitope. **A** FN-null MEFs (9.4×10^4^ cells/cm^2^) were seeded on wells pre-coated with 250 nM reduced or non-reduced S1 protein, S1-RBD, FNIII10, or 1% BSA controls. Cells were seeded for 1 h in the presence of 1 mM MnCl_2_, 10 mM EDTA, or vehicle control before cell adhesion was determined by crystal violet staining. Data are presented as mean absorbance ± SEM, n=3 independent experiemtns performed in triplicate. *p<0.05 vs corresponding BSA control by two-way ANOVA with Bonferroni’s post-hoc test. **B-E** Representative images of crystal violet-stained cells seeded in the presence of 1 mM MnCl_2_ on **B,** non-reduced S1; **C,** reduced S1; **D,** BSA; and **E,** S1-RBD. Scale bar, 100 μm.

## Discussion

The identification of a conserved RGD motif within the SARS-CoV-2 spike protein has generated substantial scientific interest (22,34), and converging lines of experimental evidence suggest that integrin inhibition may be protective against SARS-CoV-2 binding and infection (35,37–39,65). To the best of our knowledge, the data presented in the present study are the first to demonstrate that the receptor binding domain of SARS-CoV-2 spike protein functions as a classical integrin receptor agonist. S1-RBD supported cell adhesion (Fig. 1A, B) and proliferation (Fig. 1D) to comparable extents as the RGD-containing fragment of the native ECM molecule fibronectin (FNIII10). This interaction was competitively inhibited by both α_v_ integrin-blocking antibodies and RGD peptides (Fig. 2A, D), and was also observed in an SPR model of direct S1-RBD-integrin binding (Fig.3). Cells adherent to S1-RBD formed focal adhesions (Fig. 4) and key adhesion signaling mediators FAK, Src, Paxillin, and Akt were phosphorylated (Fig. 5). The present studies were conducted primarily in mouse embryonic fibroblasts that do not express detectable levels of ACE2, and thus these results are unlikely to be complicated by potential interactions of S1-RBD with ACE2. As well, S1-RBD supported both cell attachment and adhesion-based signaling in primary human small airway epithelial cells (Fig. 6). Together, these results demonstrate that SARS-CoV-2 spike protein contains a functional adhesive epitope within the RBD that mediates α_v_ integrin engagement via its RGD motif.

In contrast to the robust adhesive response observed on S1-RBD, cells seeded onto the larger S1 fragment of SARS-CoV-2 spike protein attached only weakly and exhibited limited spreading (Figs. 1C, 7B). Cell adhesion to S1 was partially rescued by chemical reduction of S1 and pretreatment of cells with Mn^2+^. One possible interpretation of these data is that the adhesive epitope contained within S1 is cryptic, and thus only available to integrins under appropriate physical and chemical conditions. DTT-sensitive, matricryptic epitopes have been identified in a number of native ECM proteins, including thrombospondin (62), which contains a cryptic RGD sequence whose exposure is regulated by cell-surface protein disulfide isomerases (66). Likewise, exposure of a DTT-sensitive, self-association epitope in fibronectin (61) can be regulated by both cell-derived mechanical force (67) and proteolytic fragmentation (61). The intact, trimeric SARS-CoV-2 spike undergoes multiple conformational changes and molecular interactions during the viral invasion process (68), including conformational flexibility of the RBD domain (69,70), as well as activation by cell-surface proteases TMPRSS2 (71) and Cathepsin L (34,72). Furthermore, the well-characterized, cell-surface spike receptor ACE2 (34,71), and more recently identified coreceptor heparin sulfate proteoglycans (73,74), can associate laterally with integrins on human cell surfaces (75–78). Thus, integrins and associated intracellular signaling partners are emerging as putative components of a larger molecular complex that is targeted during SARS-CoV-2 infection. Future elucidation of the conformational requirements and activation steps enabling functional engagement of integrin receptors with SARS-CoV-2 spike in the context of established mechanisms of viral attachment and invasion represents an open question of substantial importance.

Of reports investigating integrin-spike interactions, α_5_β_1_ has been proposed as a receptor of interest, in part due to its functional association with ACE2 (75,76), the ability of β1-selective integrin antagonists to reduce SARS-Co-V2 invasion (35,38,39), and observed α_5_β_1_ integrin-S1-RBD interactions by ELISA (35) and SPR (79) assays. Some reports have also implicated α_v_β_3_ integrins in viral entry (37,80), while others found no effects of integrin antagonists on viral invasion (34). In the present study, we compared effects of β_1_- and β_3_-integrin blocking antibodies on cell attachment to RBD using a well-characterized fibroblast cell line that expresses both functional α_5_β_1_ and α_v_β_3_ integrin receptors (41,81). Notably, FN-null MEFs do not produce fibronectin, are cultured in the absence of serum, and do not deposit other endogenous matrix molecules, including RGD-containing thrombospondin and the α_1/2_β_1_ integrin ligand collagen, into their ECM (43). FN-null MEFs adhered to S1-RBD exclusively via α_v_β_3_ integrins, with no contribution from α_5_β_1_ integrins (Fig. 2A-C). This result was confirmed using recombinant integrins and SPR, which further indicated that the affinity of the epithelial integrin, α_v_β_6_ for S1-RBD was substantially higher than that of α_v_β_3_ (Fig. 3 and Table 1). While we did not test the integrin specificity of the larger S1 fragment in the present study, recent work by Park and colleagues (65) showed that an Fc-tagged S1 fragment could support α_v_, α_4_, or β_1_-mediated adhesion depending on cell-type specific integrin expression. Thus, the possibility remains that, like fibronectin (47,48), synergistic sequences or conformational flexibility within the larger S1 domain may confer additional, dynamic integrin selectivity. The specificity and selectivity of spike and spike fragments may further be sensitive to modulators of integrin signaling, such as heparin sulfate proteoglycan co-receptors (82) or cell-surface proteases (83), both of which have been identified as factors regulating engagement of SARS-CoV-2 virions with host target cells (34,71–73).

The demonstration of an α_v_-specific, integrin agonist functionality contained within S1-RBD protein opens multiple avenues that will be critical in expanding scientific understanding of SARS-CoV-2 and therapeutic options for a global population affected by the COVID-19 pandemic. The most immediate among these, the identification of anti-integrin therapeutics that are FDA-approved or in pre-clinical trials with potential efficacy against SARS-CoV-2 infection, is already underway (84). Integrins have been implicated in the pathophysiology of numerous respiratory viruses, including human cytomegalovirus (85), hantaviruses (86), and influenza (87), as either primary receptors or as major mediators of host response and disease severity, with the specific contributions of integrins in the context of COVID-19 disease yet to be elucidated (22,88). Meanwhile, numerous questions remain unanswered regarding mechanisms underlying the differential susceptibility of vulnerable populations to severe manifestations of COVID-19 (4), as well as potential differences in infectivity, transmissibility, disease severity, and immune invasion associated with novel variants (89,90). Interrogating these open challenges in the context of integrin-spike interactions, including factors determining integrin selectivity and specificity, is a promising and yet unexplored avenue. For example, ACE2 cell surface expression levels alone do not sufficiently predict tissue susceptibility or disease severity (91). Thus, a combinatorial expression profile of ACE2, alongside α_v_ integrin surface expression may better predict cell tropism of SARS-CoV-2. Alternatively, fibronectin-integrin interactions play a key role in maintaining endothelial barrier function during sepsis (16,17,92–95), which may be disrupted by competition from spike protein fragments during SARS-CoV-2-driven inflammation. Variations in integrin expression and activation state are likewise associated with some of the key risk factors for severe complications of SARS-CoV-2 infection (4), including diabetes (96,97), hypertension (98–100), and differing inflammatory responses (87,101). As the global COVID-19 pandemic approaches a new, endemic stage, targeting the emerging spike-integrin signaling axis has the potential to become an essential tool in preventing or mitigating the most severe effects of the disease, particularly for vulnerable patients who are not fully protected by current preventative and therapeutic regimens.

## Conclusions

The SARS-CoV-2 spike protein contains a novel RGD motif within its receptor-binding domain (S1-RBD). We demonstrate that S1-RBD is a functional integrin agonist with selectivity for α_v_ integrins, specifically α_v_β_3_ and α_v_β_6_. In contrast, we found no evidence of S1-RBD engagement with α_5_β_1_ integrins in either cellular adhesion or SPR systems. S1-RBD-mediated cellular adhesion supported cell spreading and cytoskeletal engagement, focal adhesion formation, and stimulation of key intracellular signaling pathways associated with cytoskeletal organization and cell proliferation. Together, these data point to a functional role for α_v_ integrins during attachment and invasion of SARS-CoV-2, and provide insight into critical open questions regarding COVID-19 pathophysiology, including mechanisms underlying variable disease severity, intersecting risk factors, and postacute viral sequelae.

## Experimental procedures

### Reagents

Fibronectin was purified from outdated human plasma (American Red Cross, Rochester, NY) using gelatin-Sepharose (GE Life Sciences, now Cytiva) affinity chromatography (102). Type I collagen (rat tail) was purchased from Corning (354236). Unless otherwise indicated, chemicals were obtained from J.T. Baker or Sigma-Aldrich. GST-tagged FNIII10 and HN-tagged FNIII10-13 were produced and purified from *E. coli* as described previously (47,103). His-tagged S1 and S1-RBD of SARS-CoV-2 were purchased from Sino Biological (40591-V08H) and R&D Systems (10523-CV;), respectively. Where indicated, S1 was reduced by successive 1 h treatments with 10 mM DTT and 30 mM N-ethyl maleimide (NEM) at 37 °C. Both reduced and non-reduced S1 were dialyzed into PBS prior to use. Integrin-blocking antibodies anti-α_5_ (clone 5H10-27), anti-α_v_ (clone H9.2B8), anti-β_1_ (clone Ha2/5), anti-β_3_ (clone 2C9.G2), and isotype controls were purchased from BD Biosciences. Antibodies for immunofluorescent staining were as follows: vinculin (clone VIN-11-5, Sigma or clone 42H89L44, Invitrogen); phosphotyrosine (clone 4G10, Sigma or PY20, BD Biosciences); phospho-FAK pY407 (polyclonal, Invitrogen #44650G); phospho-FAK pY397 (polyclonal, Biosource #44-624G); phospho-Src pY418 (polyclonal, Biosource #44-660); phospho-Paxillin pY118 (polyclonal, Invitrogen #44-722G); phospho-Paxillin pY31 (polyclonal, Invitrogen #44-720G); phospho-Akt pS473 (polyclonal, Cell Signaling #9271); TRITC-labeled phalloidin (Millipore, #90228). Alexa Fluor-conjugated secondary antibodies were from Molecular Probes. RGD-containing peptides derived from SARS-CoV-2 (ADSFVIRGDEVRQIAPGQTG) and KGD-containing peptides derived from SARS-CoV (ADSFVVKGDDVRQIAPGQTG) were produced by Genscript. Integrin-blocking (GRGDSP, #SCP0157) and negative control (GRADSP, #SCP0156) peptides were purchased from Sigma. Recombinant human integrins αvβ3 (3050-AV), αvβ6 (3817-AV), and α5β1 (3230-A5) were from R&D Systems.

### Cell culture

FN-null MEFs were cultured under serum- and fibronectin-free conditions on collagen I-coated tissue culture flasks using a 1:1 mixture of Aim V (Invitrogen) and Corning SF Medium (Corning), as described (41). Adult human small airway epithelial cells (SAECs) were purchased from Lonza (CC-2547) and used between passage 6 and 8. SAECs were cultured in serum-free Small Airway Epithelial Growth Media (Lonza CC-3118), according to manufacturer’s instructions. Cells were passaged at 70-80% confluence using ReagentPack subculture reagents (Lonza CC-5034). Neither FN-null MEFs nor SAECs expressed detectable levels of ACE2 protein by immunoblot blot analysis (data not shown).

### Cell adhesion and proliferation assays

Cell adhesion assays were performed as described previously (81). Briefly, 96-well tissue culture plates were coated with S1-RBD (10 - 1000 nM), FNIII10 (10-1000 nM), GST (1000 nM), or S1 (7.8 - 250 nM) for 1 h at 37 °C. Cells were seeded on protein-coated wells (9.4 x 10^4^ cells/cm^2^) in either AimV/SF medium (FN-null MEFs) or Small Airway Epithelial Basal Medium (CC-3119; Lonza) in the absence or presence of EDTA (10 mM), DTT (1 mM), or MnCl_2_ (1 mM) as indicated; MnCl_2_ was added 1 h after seeding. For integrin blocking studies, FN-null MEFs were pre-incubated with anti-integrin antibodies (50 μg/mL) or 25 μM peptide for 1 h prior to seeding. Cells were then seeded into wells and incubated at 37 °C and either 8% (Fn-null MEFs) or 5% (SAECs) CO_2_ for up to 2 h. Wells were then washed with PBS to remove non-adherent cells, fixed with 1% paraformaldehyde and stained with 0.5% crystal violet. The absorbance of crystal violet solubilized in 1% SDS was measured at 590 nm. Proliferation assays were performed by seeding FN-null MEFs (2.5 x 10^3^ cells/cm^2^) on protein-coated 48-well plates. Cells were cultured for 4 days at 37 °C, 8% CO_2_ and then fixed and stained with crystal violet (81). In some experiments, images of adherent cells were obtained after crystal violet staining and before solubilization, using an IX70 inverted microscope (Olympus) equipped with a Micropublisher 3.3 RTV digital camera (Q Imaging).

### Surface plasmon resonance

Kinetic studies of integrin-ligand interactions were performed using a BIAcore T200 instrument (Cytiva). Ligands (S1-RBD or FNIII10) were immobilized using amine-coupling chemistry according to the manufacturer’s instructions (BR-1000-50). Briefly, ligands diluted in 10 mM sodium acetate (pH 4.0, Cytiva) were immobilized on an EDC/NHS-activated CM5 chip (Cytiva) to a target level of 800-1000 RU. Excess amine-reactive groups were inactivated with 1 M ethanolamine (pH 8.5, Cytiva). Immobilization buffer was 10 mM HEPES buffer pH 7.4 containing 0.05% n-octyl-β-D-glucopyranoside (OGPS, Anatrace), 150 mM NaCl_2_, 2 mM MnCl_2_, 2 mM MgCl_2_, and 0.2 mM CaCl_2_. Lyophilized integrins were reconstituted with 50 mM Tris, pH 7.4 containing 25 mM OGPS, 1 mM DTT, 150 mM NaCl_2_, and divalent cations (α_v_ integrins: 2 mM MnCl_2_, 2 mM MgCl_2_, and 0.5 mM CaCl_2_; α_5_β_1_ integrin: 2 mM MnCl_2_). Double-referenced binding experiments were performed in parallel flow cells for S1-RBD and the corresponding positive control (FNIII10 for α_v_ integrins; FNIII8-10 for α_5_β_1_)(47) using a flow rate of 30 μL/min for 1 min with a dissociation time of 5 min. Surfaces were regenerated between injections using two 30 s injections of 20 mM EDTA and 1M NaCl (48). Kinetic parameters were determined by fitting a 1:1 binding model with globally fit parameters for each collected data set using Biacore T200 Evaluation software (Version 3.2, GE). Due to the large difference in dissociation rates between the two ligands, only the first 10 seconds of the dissociation curves were considered for S1-RBD data sets. Quality of fit was determined by agreement between measured and calculated Rmax and Chi-squared values. Data sets not producing high-quality kinetic fit were excluded from calculation of kinetic parameters.

### Immunofluorescence microscopy

Acid-washed glass coverslips were coated with 500 nM protein (S1-RBD or FNIII10) for 1 h at 37 °C. FN-null MEFs (2.5×10^3^ cells/cm^2^) were seeded in AimV/SF media and incubated at 37 °C, 8% CO_2_ for 4h. Cells were then fixed with 2% paraformaldehyde in PBS and processed for immunofluorescence microscopy as described previously (104). Cells were incubated with primary antibodies or TRITC-phalloidin diluted in PBS containing 0.1% Tween 20, 1% BSA and 1 mM phenylmethylsulfonyl fluoride for 1 h at room temperature. Bound antibodies were detected with Alexa^448^-, Alexa^549^-, or Alexa^647^-labeled goat anti-rabbit or -mouse secondary antibodies and visualized using a BX60 fluorescence microscope (Olympus) equipped with an epifluorescent lamp (Lumen Dynamics) and an EXi Blue Fluorescence Camera (Q Imaging), acquired with QCapture software.

### Immunoblot analysis

FN-null MEFs (3.4 x 10^4^ cells/cm^2^) or SAECs (6.7 x 10^4^ cells/cm^2^) were seeded on wells pre-coated with S1-RBD (500 nM), FNIII10 (500 nM), or fibronectin (10 μg/ml) and incubated at 37 °C for 1 h (FN-null MEFs) or 4 h (SAECs). Cells were lysed with 40 μL/cm^2^ SDS-RIPA buffer (50 mM Tris, 150 mM NaCl, 1 mM EDTA, 1% Triton X-100, 0.1% sodium dodecyl sulfate, 0.5% sodium deoxycholate, pH 7.6 containing 1 mM sodium orthovanadate, 1 mM phenylmethylsulfonyl fluoride, and 1X protease inhibitor cocktail (Sigma S8830). Cell lysates were analyzed by SDS-PAGE and immunoblotting (105). Immunoblots were blocked with either 5% non-fat milk or 3% BSA in Tris-buffered saline containing 0.1% Tween 20 (TBS-T). Membranes were incubated overnight at 4 °C with primary antibodie diluted in TBS-T. Blots were then washed with TBS-T, incubated with horseradish peroxidase-conjugated secondary antibodies, and developed using SuperSignal™ West Pico Chemiluminescent Substrate (Thermo Scientific). Blots were imaged using a ChemiDoc imaging system (Bio-Rad).

### Statistical analysis

Data are presented as mean ± standard error unless otherwise stated. Experiments were performed in triplicate on a minimum of 3 independent days. All statistical analyses were performed using GraphPad Prism (version 9). Statistical differences between groups were identified by one- or two-way ANOVA as indicated, using Bonferroni’s post-test and a p-value threshold <0.05.

## Data availability

All data are contained within the manuscript, with supplementary data available upon request

## Acknowledgements

The authors thank the University of Rochester Structural Biology and Biophysics Facility for the use of the Biacore T200 System, and Dr. Jermaine L. Jenkins for technical assistance.

## Author Contributions

E. G. N. and D. C. H. conceptualization; E. G. N. and D. C. H. data curation; E. G. N. and X. S. P. formal analysis; E. G. N., X. S. P. and D. C. H. investigation; E. G. N. writing-original draft; E. G. N. and D. C. H. writing - review and editing; D. C. H. supervision; D. C. H. funding acquisition; D. C. H. methodology; D. C. H. project administration

## Funding Sources and Additional Information

This research was supported by National Institutes of Health grant R01 AG058746 and by a University Research Award from the University of Rochester

